## Supplementary Information for "Structural basis for specific RNA recognition by the alternative splicing factor RBM5"

#### Figure S1

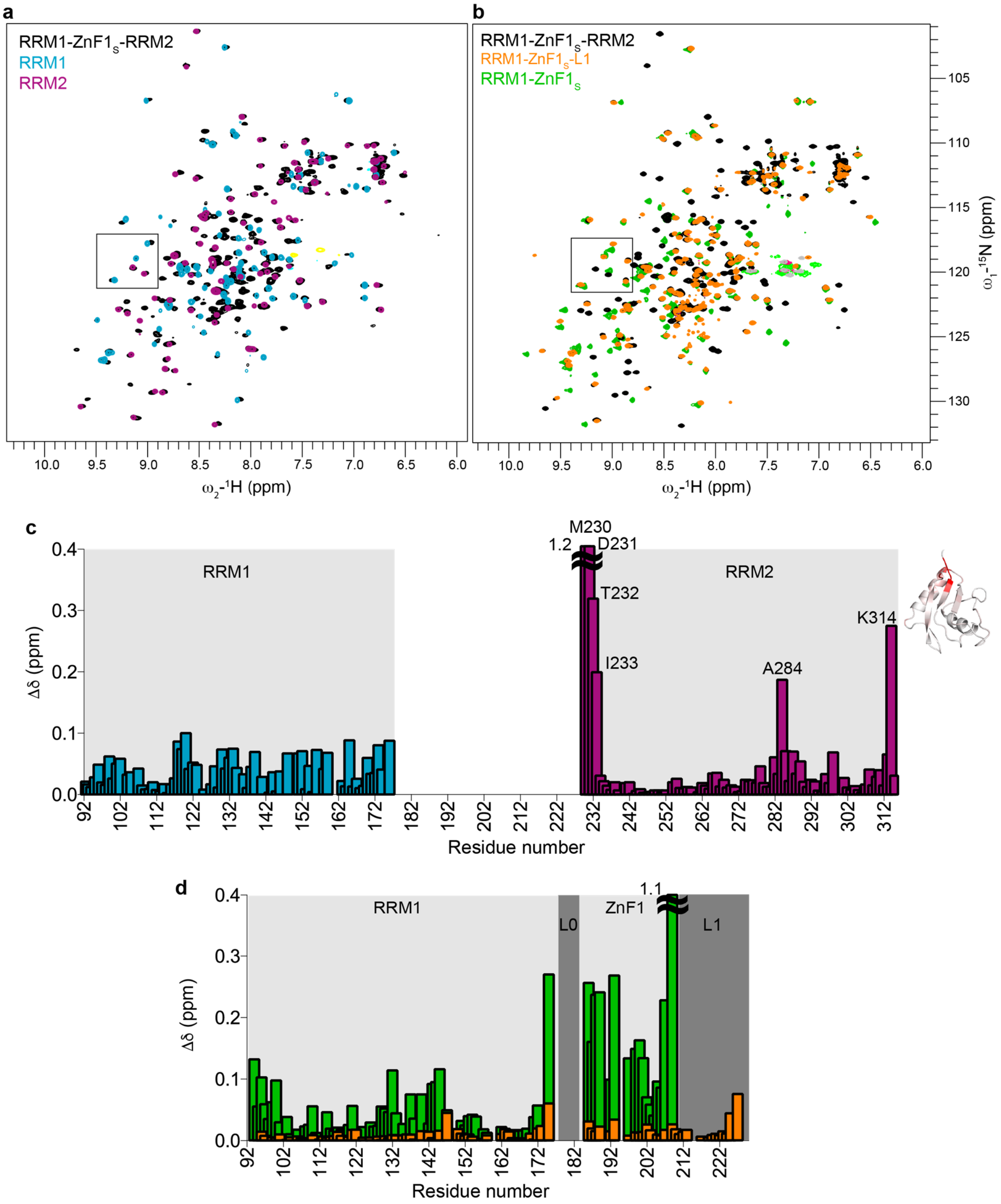

**Figure S1. $^{1}H$-$^{15}N$ HSQC spectra of single and multidomain constructs of RBM5 RNA binding domains.**

**(a)** Overlay of $^{1}H$-$^{15}N$ HSQC spectra of three RRM1-ZnF1$_S$-RRM2 domain construct (black) and single RRM1 (light blue) and RRM2 (purple) domains. Zoomed view of residues shown in **Fig. 1c** is marked. The spectra were recorded in buffer containing 20 mM MES pH 6.5, 100 mM NaCl, 1 mM DTT. **(b)** Overlay of $^{1}H$-$^{15}N$-HSQC spectra of the three RRM1-ZnF1$_S$-RRM2 domain construct (black) with that of RRM1-ZnF1$_S$-L1 (orange) and RRM1-ZnF1$_S$ (green). Zoomed views shown in **Fig. 1d** are indicated by rectangles. NMR spectra were recorded in buffer containing 20 mM MES pH 6.5, 400 mM NaCl, 1 mM DTT. **(c)** Chemical shift differences for amide signals in RRM1 (light blue) and RRM2 (purple) compared to three RRM1-ZnF1$_S$-RRM2 domains (corresponding to $^{1}H$-$^{15}N$ HSQC spectra in panel a). For RRM2, the chemical shift differences are plotted onto the structure (PDB ID: 2LKZ) in a white to red colour scheme. **(d)** Chemical shift differences of amide signals in RRM1-ZnF1$_S$ (green) and RRM1-ZnF1$_S$-L1 (orange) compared to the three RRM1-ZnF1$_S$-RRM2 domains (corresponding to $^{1}H$-$^{15}N$ HSQC spectra in panel b).

#### Figure S2

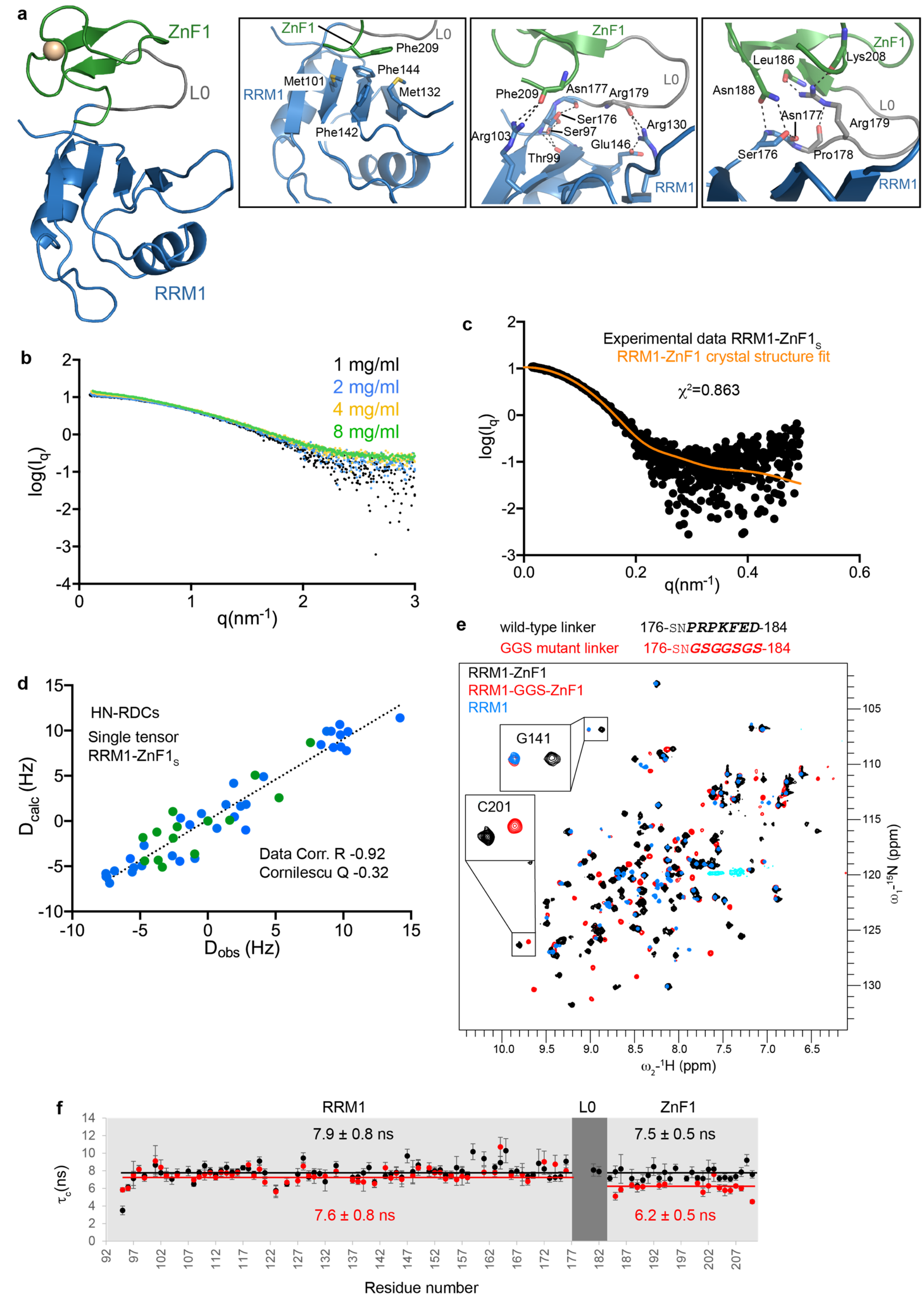

**Figure S2. RRM1-ZnF1 crystal structure and solution conformation.**

**(a)** The crystal structure of RRM1-ZnF1 tandem domains is shown with RRM1, ZnF1, linker L0 and $Zn^{2+}$ ion colored in blue, green, gray and golden, respectively. The structure shows a tri-partite mode of interaction between

RRM1, ZnF1 and the linker connecting them (L0). Zoomed views of the interaction interface are shown in boxed panels. Dotted lines represent hydrogen bonds. **(b-f)** Validation of the solution conformation of RRM1-ZnF1. Due to instability of RRM1-ZnF1 at room temperature for longer durations, the C191G mutant was used for SAXS and NMR based RDC experiments. **(b)** Concentration dependent increase in intensity of scattering profiles is observed in RRM1-ZnF1$_S$ protein. **(c)** The fit between experimental SAXS data for RRM1-ZnF1$_S$ protein at 1 mg/ml vs. data back-calculated from the RRM1-ZnF1 crystal structure is shown (using Crysol), the $\chi^2$ value is indicated. **(d)** Analysis of $^1$H-$^{15}$N RDCs (using 3% PEG-hexanol alignment medium) is shown. The experimental or observed RDCs ($D_{obs}$) are plotted against the back-calculated RDCs ($D_{calc}$) from the crystal structure and the data correlation $R$ factor and Cornilescu $Q$ factor are indicated. **(e)** To investigate the importance of the linker L0 as an anchor between the two domains, residues 178-184 were replaced by a stretch of Gly-Gly-Ser repeats of the same length (RRM1-GGS-ZnF1). Since many of the contacts of the linker with either of the domains are mediated by side-chain interactions, this mutant disrupts inter-domain contacts. Superposition of wild-type RRM1-ZnF1 (black), RRM1-GGS-ZnF1 linker mutant (red) and RRM1 (blue) single domain $^1$H-$^{15}$N HSQC spectra show very large chemical shift differences indicating a significant structural change. All data were collected in low salt buffer (100 mM NaCl) and at low concentrations for tandem domain constructs (due to instability of the protein in low salt conditions are high concentrations). Zoomed views of two residues (Gly141, Cys201) demonstrate that the chemical shifts of residues in single domains are comparable to those in the RRM1-GGS-ZnF1 linker mutant **(f)** Local rotational correlation time ($\tau_c$) are calculated from $^{15}$N $R_1$ and $R_2$ relaxation rates *vs.* residue numbers for wild-type RRM1-ZnF1 (black), RRM1-GGS-ZnF1 linker mutant (red). The average and standard deviation is listed for each domain. The faster tumbling of the ZnF1 domain with a reduction of $\tau_c$ from 7.5 ns to 6.2 ns from wild-type to the RRM1-GGS-ZnF1 mutant indicates that the linker mutation disrupts interactions between the linker and the two flanking domains. The flexible GGS-linker allows the two domains to tumble independently, with domain-specific tumbling correlation times that reflect their size difference.

#### Figure S3

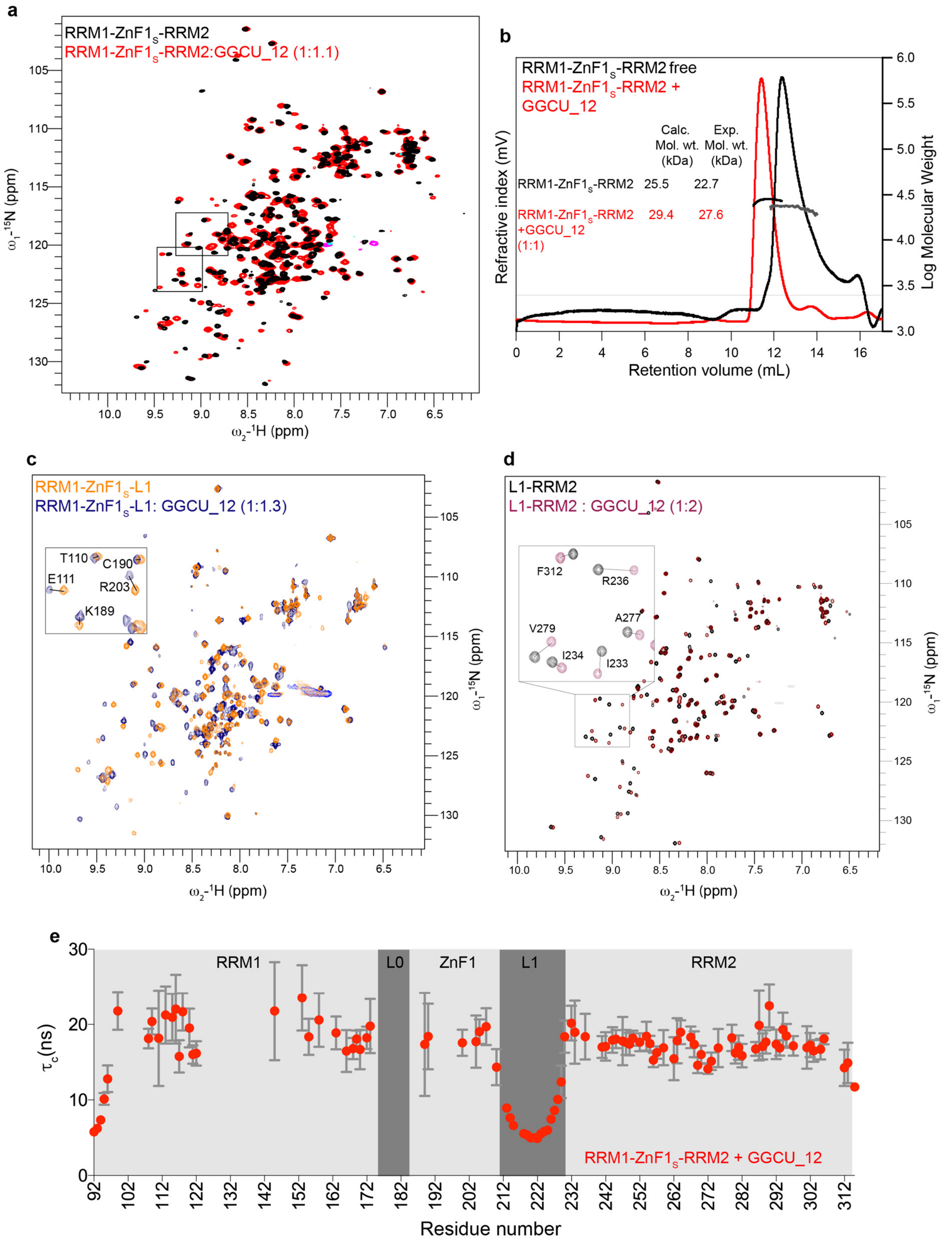

**Figure S3. RRM1, ZnF1 and RRM2 all bind GGCU_12**

**(a)** Overlay of $^{1}$H-$^{15}$N HSQC spectra of RRM1-ZnF1$_S$-RRM2 free (black) and bound to GGCU_12 RNA (red). Zoomed views of residues shown in **Fig. 2a** are marked, the corresponding CSP plot is shown in **Fig. 2b**. **(b)** Size-exclusion chromatography coupled to static light scattering for measurement of molecular weights of RRM1-ZnF1$_S$-RRM2 free (black) and in complex with GGCU_12 (red). The peak regions used for calculation of experimental molecular weights are marked. The calculated molecular weight from sequence (Calc. Mol. wt.) and experimental molecular weights (Exp. Mol. wt.) are indicated. **(c)** Overlay of $^{1}$H-$^{15}$N HSQC spectra of RRM1-

ZnF1$_S$-L1 -/+ GGCU_12 (orange, blue), corresponding to CSP plot in **Fig. 2b**. **(d)** Overlay of $^{1}$H-$^{15}$N HSQC spectra of L1-RRM2 -/+ GGCU_12 (black, purple), corresponding to CSP plot in **Fig. 2b**. **(e)** Linker L1 remains flexible in RRM1-ZnF1$_S$-RRM2 upon RNA binding. Local tumbling rotational correlation time ($\tau_c$) calculated from $^{15}$N $R_1$ and $R_2$ rates *vs.* residue number for RRM1-ZnF1$_S$-RRM2 + GGCU_12 RNA (red).

#### Figure S4

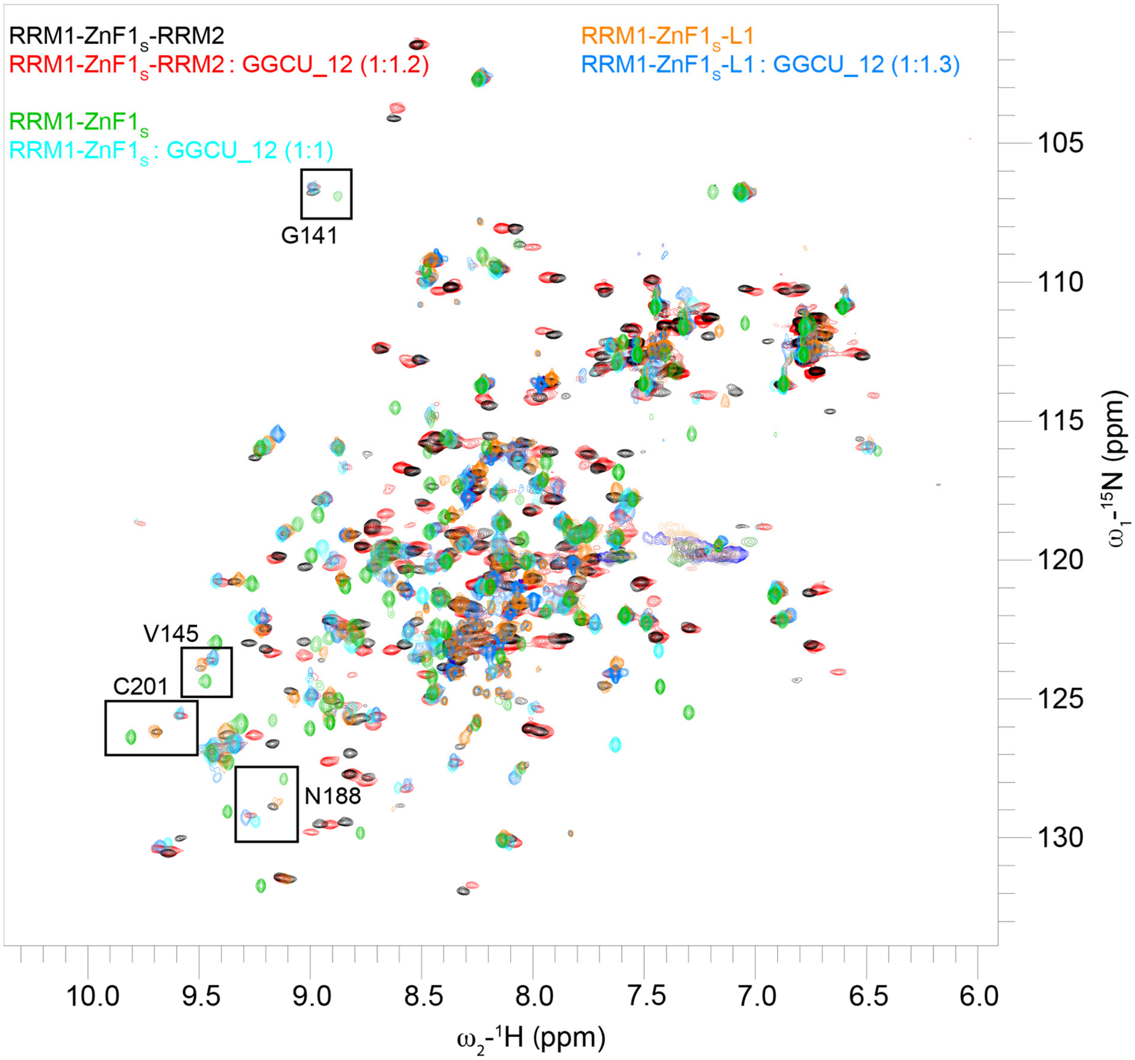

**Figure S4. RNA binding by RRM-ZnF1$_S$-RRM2, RRM1-ZnF1$_S$-L1 and RRM1-ZnF1$_S$ is similar.** Overlay of $^{1}$H-$^{15}$N HSQC spectra of RRM1-ZnF1$_S$-RRM2 -/+ GGCU_12 (black, red), RRM1-ZnF1 $_S$-L1 -/+ GGCU_12 (orange, blue) and RRM1-ZnF1$_S$ -/+ GGCU_12 (green, cyan) are shown. Zoomed views of residues shown in **Fig. 2d** are marked.

### Figure S5

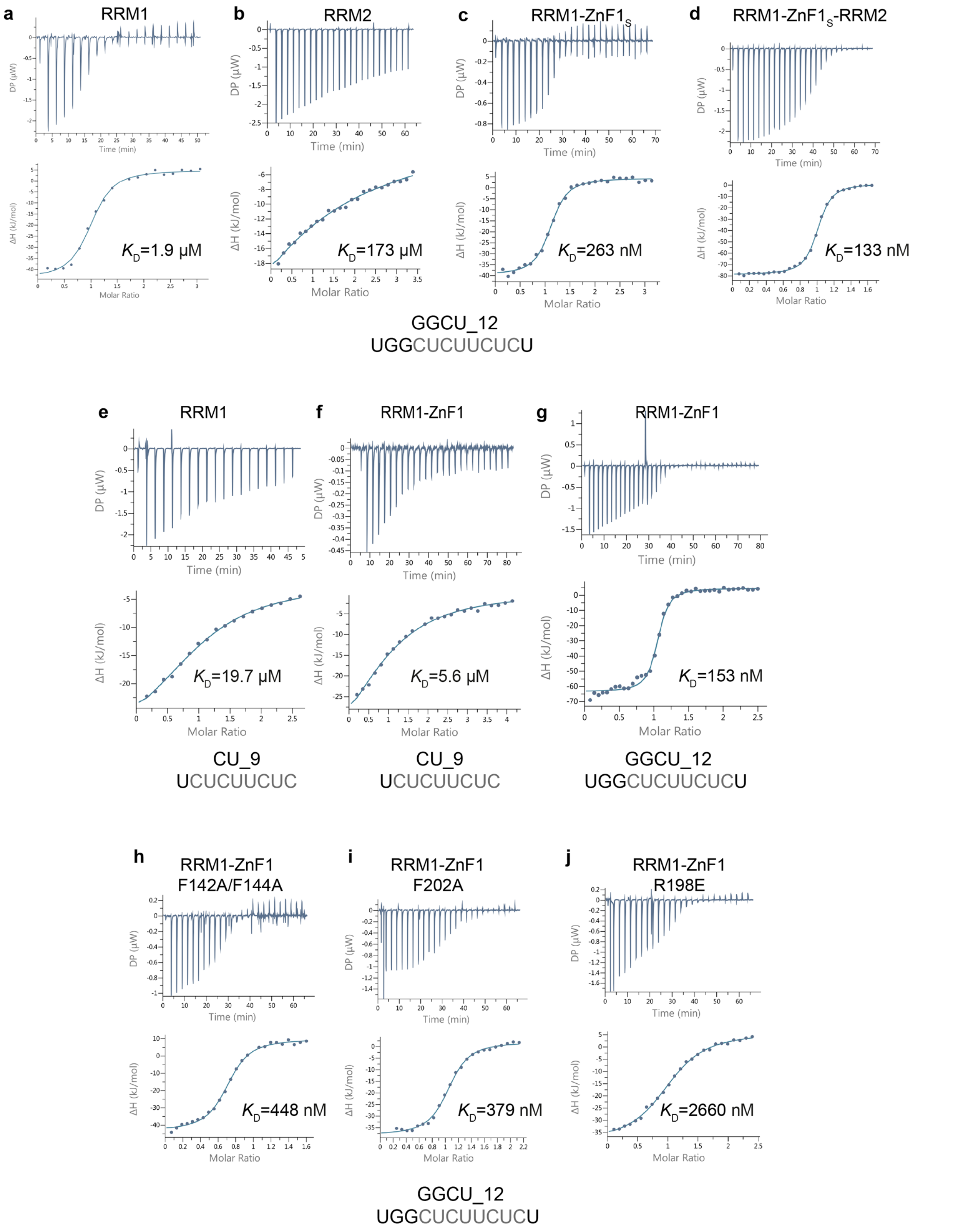

**Figure S5. ITC binding isotherms.**

**(a-d) Cooperative binding by the different RNA binding domains of RBM5.** ITC binding isotherms of RRM1, RRM2, RRM1-ZnF1$_S$ and RRM1-ZnF1$_S$-RRM2 with GGCU_12 are shown in panels a-d, respectively. **(e-g)** RRM1 and ZnF1 both have RNA binding affinity contribution towards the GG motif. ITC binding isotherms of RRM1 and RRM1-ZnF1 with CU_9 and RRM1-ZnF1 with GGCU_12 are shown in panels e-f, respectively. **(h-j)** RNA binding contribution of specific residues of RRM1 and ZnF1. ITC binding isotherms of RRM1-ZnF1$_{F142A/144A}$, RRM1-ZnF1$_{F202A}$ and RRM1-ZnF1$_{R198E}$ with GGCU_12 are shown in panels h-j, respectively. The binding affinities derived from duplicate measurements are listed.

#### Figure S6

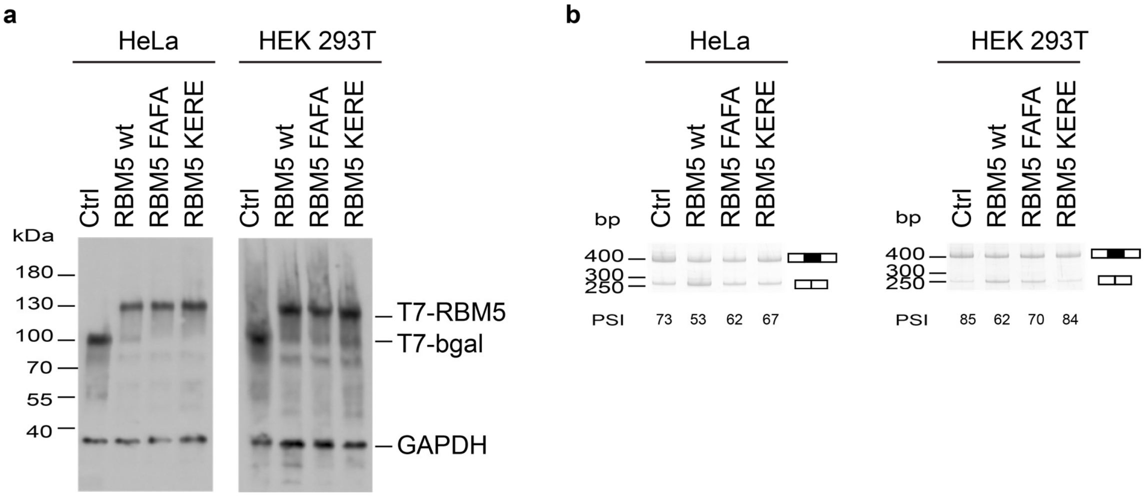

**Figure S6. Activity of RBM5 RRM1 and ZnF1 mutants**

HeLa and HEK 293T cells were co-transfected with RG6 NUMB alternative splicing reporter and T7-RBM5 vectors expressing wild type or RNA binding affinity mutants in the RRM1 (F142A/F144A-> FAFA) or ZnF1 (K197E/R197E-> KERE) or control vector (expressing beta-galactosidase), as indicated. Protein expression was assessed by western blot using anti-T7 epitope antibodies and protein loading was assessed by measuring the levels of GAPDH. **(a)** Pattern of alternative splicing isoforms was detected by RT-PCR using primers complementary to vector sequences flanking exons of the RG6-NUMB minigene; the positions of the amplification products corresponding to exon 9 inclusion/skipping are indicated. **(b)** The results correspond to one representative replicate of the experiment. Quantification of alternatively spliced isoforms from multiple replicates such as in (b) were used to generate the histogram shown in **Fig. 4d**.

### Figure S7

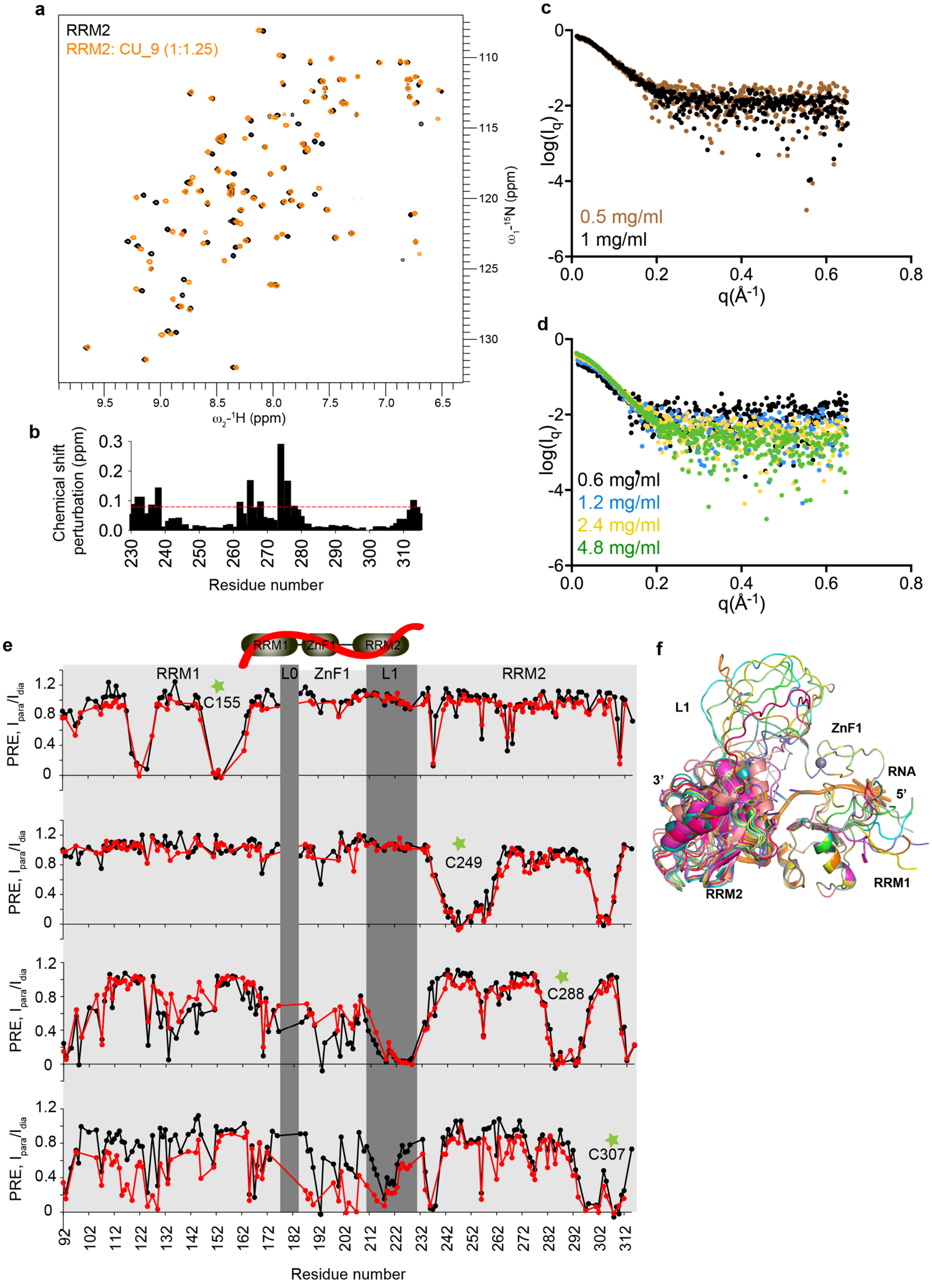

**Figure S7. Structural model of RBM5 RRM1-ZnF1-RRM2 when bound to RNA.**

**(a)** Overlay of $^{1}H$-$^{15}N$ HSQC spectra of RRM2 free (black) and bound to CU_9 RNA (orange). **(b)** Chemical shift perturbations in RRM2 upon binding to CU_9 RNA at a ratio of 1:1.25 (protein:RNA) *vs.* residue number are shown. Protein residues which satisfy a threshold RNA binding criterion ($\Delta\delta > 0.08$, shown as red dashed line):

Asp231, Thr232, Ile233, Arg236, Ile238, Ile262, Ile265, Lys268, Arg274, Phe276, Phe278 and Ala313 were used to generate ambiguous distance restraints to the RNA during structure modeling. **(c)** Small angle X-ray scattering (SAXS) profiles for RRM1-ZnF1$_S$-RRM2 at 0.5 mg/ml and 1 mg/ml. **(d)** Concentration-dependent increase in intensity of the scattering profiles is observed for the RRM1-ZnF1$_S$-RRM2 + GGCU_12 RNA complex. Data at lowest concentration were carefully merged with those at higher concentrations and used for further analysis. **(e)** PRE ratio of NMR signal intensities in the paramagnetic and diamagnetic state, ($I_{para}/I_{dia}$) observed in RRM1-ZnF1$_S$-RRM2 free (blue) and bound to GGCU_12 RNA (grey) *vs.* residue number. Spin label positions are marked by green stars: C155, C249, C288 and C307. **(f)** Superposition of 10 lowest energy structures of RRM1-ZnF1$_S$-RRM2 in the presence of RNA obtained from PRE-based structure calculations is shown in different colours.

#### Figure S8

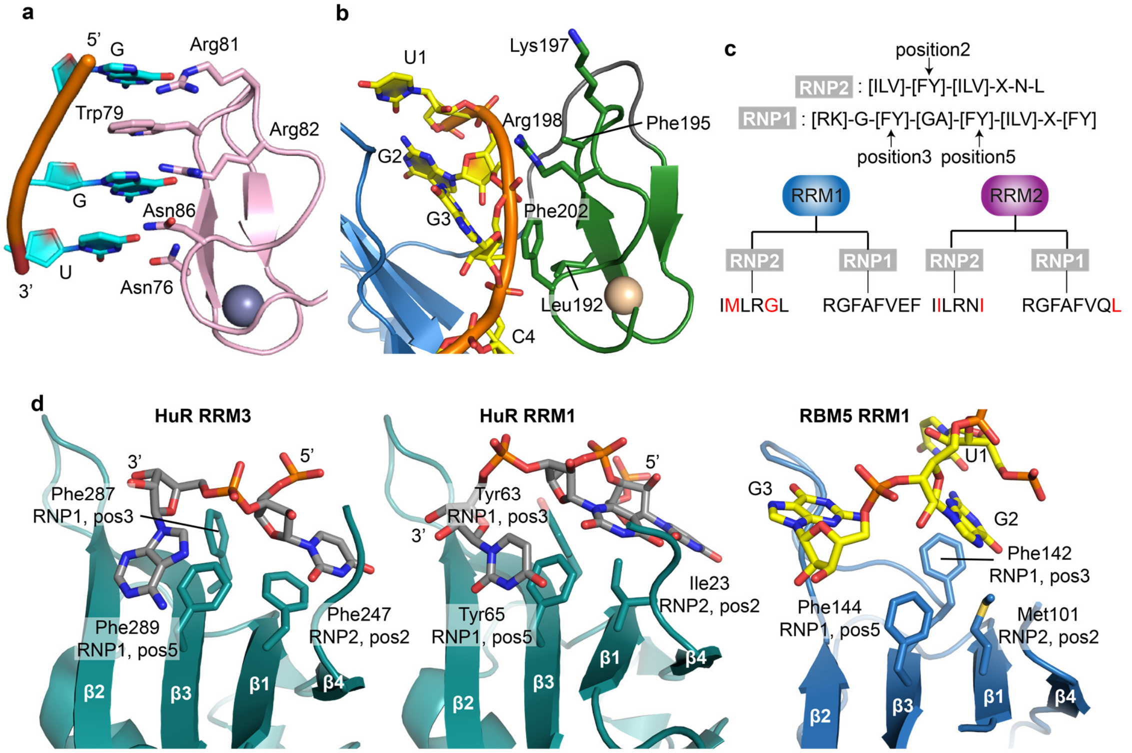

**Figure S8. Comparison of RBM5 RRM1-ZnF1 with canonical RanBP2-type zinc finger and RRMs**

**(a)** Structure of RanBP2-type Zinc finger domain (ZRANB2-F2) in complex with GGU motif RNA (PDB ID: 3G9Y[1]) is shown. Gua-Trp-Gua ladder formed between the protein and RNA is indicated. **(b)** A view of RBM5 RRM1-ZnF1$_S$ bound to GGCU_10 RNA crystal structure. Residues corresponding to ZRANB2 F2 involved in RNA recognition are marked. **(c)** Canonical RNP2/RNP1 residues of RRM domains are represented, with those in RBM5 RRM1 and RRM2 deviating from the canonical sequences shown in red. **(d)** Crystals structures of HuR RRM3 (teal) bound to RNA (grey) (PDB ID: 6GD3[2]), HuR RRM1 (teal) bound to RNA (grey) (PDB ID: 4ED5[3]) and RBM5 RRM1 (blue) bound to RNA (yellow) (this study, linker L0 and ZnF1 are not shown). For simplicity, only 2-3 nucleotides stacking with position2 of RNP2, positions 3 and 5 of RNP1 of the different RRMs are shown.

### Figure S9

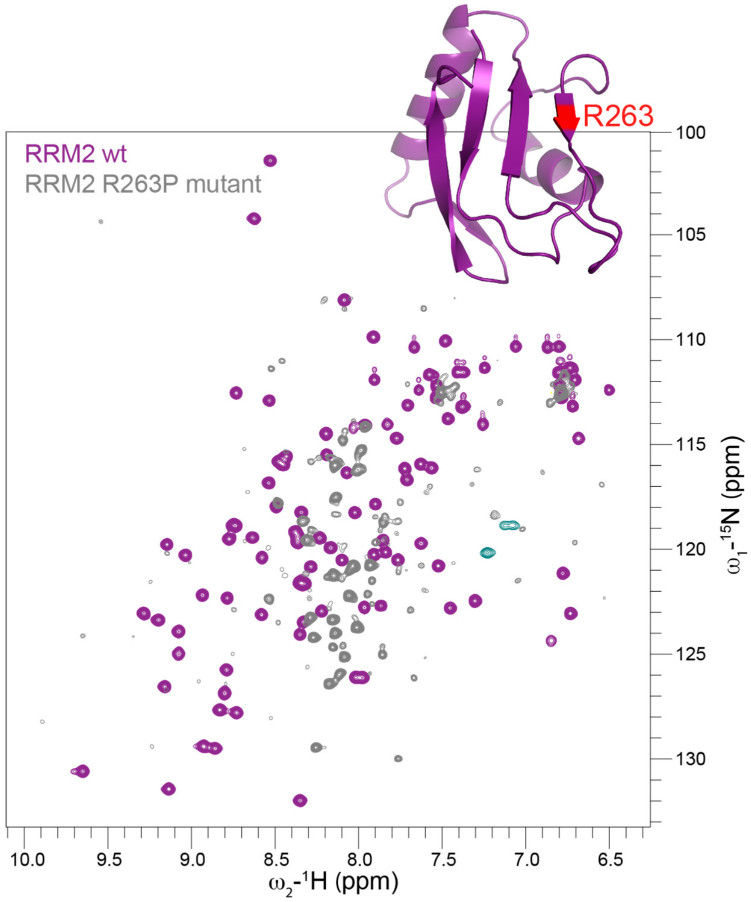

**Figure S9. Male sterility associated mutation in RBM5 RRM2**

A superposition of $^{1}H,^{15}N$ -HQSC spectra of wild-type RRM2 (purple) and RRM2 R263P mutant protein (grey) is presented. The position of point mutation is shown on the structure of RRM2 (PDB ID: 2LKZ) in red. R263P compromises the structural integrity of RBM5 RRM2.

#### Table S1

| Parameters | RRM1-ZnF1$_S$ | RRM1-ZnF1$_S$-RRM2 apo | RRM1-ZnF1$_S$-RRM2: +GGCU_12 RNA |
|---|---|---|---|
| **Data-collection** | | | |
| Mode | Batch | Batch | Batch |
| Instrument | BioSAXS BM29 ESRF | Rigaku BIOSAXS1000 | Rigaku BIOSAXS1000 |
| Beam geometry | 10 mm slit | 10 mm slit | 10 mm slit |
| Wavelength (Å) | 0.9919 | 1.5 | 1.5 |
| q range (Å$^{-1}$) | 0.0029-0.494 | 0.004-0.65 | 0.004-0.65 |
| Exposure time (s) | 15 (15x1s) | 7200 (8x900s) | 7200 (8x900s) |
| Concentration (mg/ml) | 1 | 1 | Data merged (unknown from SEC) |
| Temperature (°C) | 20 | 5 | 5 |
| **Structural parameters** | | | |
| $R_g$ (Å) [from p(r)] | 16.822 | 24.3 | 22.56 |
| $R_g$ (Å) [from Guinier] | 16.7 ± 0.02 | 23.3 ± 1.93 | 22.29 ± 3.52 |
| $D_{max}$ (Å) | 56 | 78 | 75 |
| **Molecular weight determination (kDa)** | | | |
| From Size & Shape | 15.0 | 23.9 | 27.7 |
| From volume of correlation ($V_c$) | 14.9 | 19.7 | 24.7 |
| Calculated *MW* from sequence | 13.9 | 25.5 | 29.4 |
| **Software employed** | | | |
| Primary data reduction | BsxCuBE | Rigaku SAXSLab v3.0.lr1 | Rigaku SAXSLab v3.0.lr1 |
| Data processing | PRIMUS | PRIMUS | PRIMUS |
| Computation of model intensities | CRYSOL | | CRYSOL |

**Table S1. SAXS data collection and processing statistics**
